# Potential Opportunities of Modeling Bioavailability for Monoclonal Antibodies: An Overview of mAbs and the current challenges of mAb development

**DOI:** 10.1101/2024.04.14.589447

**Authors:** Aryan Kohli, Omar Fayaz, Alexandra Y. Jensen, Christian Chung, Christopher Korban

## Abstract

With a growing market size, and a large variety of applications, monoclonal antibody technology adoption and clinical usage is at an all-time high. This review article seeks to explore 10 monoclonal antibodies (mAbs) and their mechanism of action, specifically their pharmacodynamic (PD) and pharmacokinetic (PK) properties, and use a machine learning model with various parameters to assess whether the mAb has adequate bioavailability when delivered subcutaneously. This is an investigation of drug optimization and patient outcomes when transitioning from traditional IV administrations to subcutaneous injections. The machine learning model is an extension based on a paper by Han Lou and Michael Hageman, *Machine Learning Attempts for Predicting Human Subcutaneous Bioavailability of Monoclonal Antibodies*, where they took 10 mAbs and analyzed 45 different features. To further extend this paper, we took an additional 10 monoclonal antibodies that were delivered subcutaneously, and took into account their dosage concentration as an extension to traditional PK properties. By including additional mAbs and dosage, a more sophisticated model can be produced with high scalability to deep learning modalities.

## 1 Introduction

In the immune system, white blood cells, more specifically B cells, are primarily responsible for producing natural antibodies in our body to fight a variety of diseases and infections in our bodies. There are two types of B cells including plasma cells and memory b-cells both with different functionalities. When a foreign particle enters the body, the plasma cells can rapidly produce antibodies to fight the infection. The memory cells can remember the antibody so that if the same foreign antigen were to be internalized to the body, it would be able to generate a response against the antigen. While natural antibodies develop responses to foreign antigens, there may be reasons as to why monoclonal antibodies (mAbs) could be a preferred treatment for a disease and infection. Oftentimes, the natural antibodies can produce a polyclonal response that could be used to fight a variety of antigens and can react with other proteins than the intended target. Monoclonal antibodies on the other hand are designed to target a specific antigen protein. This can be highly valuable in therapies for specific diseases that involve a specific type of antigen that can be targeted by artificially synthesized monoclonal antibody drugs.

Monoclonal Antibodies are derived from hybridoma cells which are a combination of B-lymphocyte cells and myeloma cells. The B-lymphocyte, which is responsible for creating antibodies, can come from mice (murine), humans, or both (chimeric). To formulate the B-lymphocyte, the antigen of interest is injected into the animal/human to produce the B-lymphocyte cells. From there, it is fused with myeloma cells to produce a large supply of hybridoma cells in culture. Without the myeloma cells, the B-lymphocytes would not be able to survive in culture. Once the hybridoma cells can mature in culture, they can be cloned to produce many of the same monoclonal antibodies. This was first discovered in 1975 by Georges Köhler and César Milstein. 11 years after this discovery in 1986, the FDA approved the first monoclonal antibody drug Orthoclone OKT3 (muromonab-CD3).

A variety of diseases can be treated with mAbs including chronic inflammatory conditions, the inhibition of angiogenesis in cancers, systemic lupus erythematosus, melanoma, multiple sclerosis, breast cancer, Crohn’s disease, and many more. Because of the wide variety of applications, there has been a rapid growth in these immunotherapies. By 2025, there is expected to be 300 billion dollars in revenue in the mAbs market [1]. In addition, the compound annual growth rate for the global market is at 11.07% with an expected market size of 679 billion dollars by 2033.

## 2 Methods

**Figure 1:**
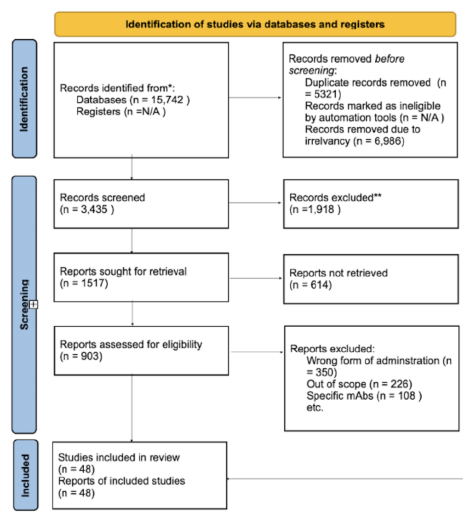
A systematic approach to review with methods following PRISMA guidelines. Reports excluded were due to irrelevance to our study, wrong form of administration, or out-of-scope interventions. The databases that were used include Drugbank, Pubmed, and Google Scholar. Additional articles demonstrating market trends for specific mAb drugs were also used in the scope of this review article.

### 2.1 Methodology

The methodology for this review on monoclonal antibodies followed the PRISMA 2020 guidelines. To find literature for the design of mAbs, subcutaneous administration, pharmacodynamics, and pharmacokinetics of specific mAbs as well as their bioavailability that covered a wide range of diseases. The collection of data and information came from a wide variety of established peer-reviewed publications from the databases PubMed and Google Scholar. Specific terms were used in the search such as “Monoclonal Antibodies”, “subcutaneous administration”, “pharmacokinetics”, and “bioavailability” to properly tailor the research process. Manual research regarding market analysis of certain mAbs drugs was also a part of the research process. Publications excluded from this research were due to the irrelevant scope of the topic, wrong form of administration, and mAbs drugs that were not included in our review analysis. The initial screening process included 15,742 articles. Next, duplicated articles (n = 5321) and irrelevant articles (n = 6986) were removed from the search. Additional articles were excluded because they did not pertain to the scope of the review article (n=1918). With 1517 articles eligible to be screened, 614 publications were not able to be retrieved due to limited access to full text. Lastly, a more detailed search was applied excluding non-subcutaneous administration and limited pharmacokinetic and pharmacodynamic trial data. A total of 40 articles were included in the review that encompassed the detailed structure of monoclonal antibodies, specific types of mAbs, the pharmacokinetic and pharmacodynamics properties, and the bioavailability of subcutaneous administration. The assessment of study quality was conducted meticulously, employing standardized assessment tools. Factors such as study design, methodological robustness, and the relevance and reliability of data were carefully considered. Moreover, adherence to ethical standards in data handling was paramount, ensuring compliance with ethical research protocols. Sensitive information was treated with confidentiality and utmost respect throughout the review process.

### 2.2 Data Collection

Antibody sequences were determined using online tools indicated in the paper, including DrugBank and Kegg drug databases. Using the amino acid sequences as well as protein data bank files, we were able to extract the following features using different software:

**Figure 2:**
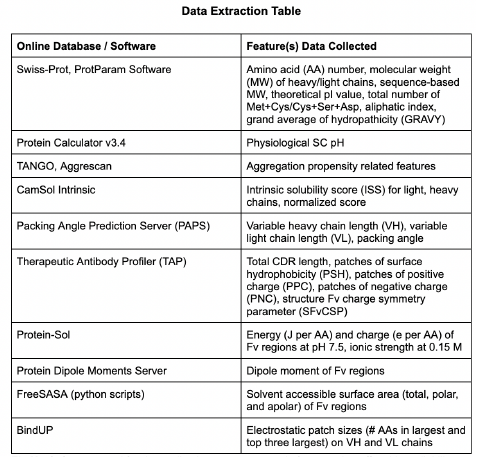
Softwares used for data collection to extract certain features that affect bioavailability

Fv region = variable domain, important for binding to antigen Formulation concentration was found in referenced clinical research papers, any concentrations that were listed in mg/kg were multiplied by 62 kilograms (average human body weight) to produce a value in mg. Formulation pH was calculated by subtracting 2.12 units (average deviation from pI) from the pI value In total, we collected 45 different parameters that contribute to the bioavailability of 23 monoclonal antibodies in our dataset.

### 2.3 Code Implementation

Using the pandas library, we imported our dataset (CSV file) and with the dataframe functions, we separated our data into an X dataset (monoclonal antibodies and features) and Y dataset (bioavailability). We normalized the data to avoid any potential scaling issues when doing principal component analysis (PCA). Our next step was to perform PCA on the dataset to identify the most relevant features and reduce dimensionality while still explaining the majority of the variance in our model. Once we performed PCA analysis and measured how much variance was used, we created a PC plot for the first two components that accounted for the most variance out of the PCs. In addition, we created a heatmap of the PC components and features from the data set to see how each feature was affecting each principal component. After finishing PCA, we had to decide which predictive model to use for the dataset. Since the model should determine whether a monoclonal antibody had adequate bioavailability, we chose a binary classification model. Because the dataset was highly dimensional, and since there was obscurity regarding whether it was linear or nonlinear separable because the hyperplane margins did not clearly separate the features, Support Vector Machine (SVM) was an appropriate technique to use. Because we are using SVM as a binary classification method, we had to convert our Y matrix (bioavailability %) to an array of 1s and 0s. Similar to the paper Machine Learning Attempts for Predicting Human Sub-cutaneous Bioavailability of Monoclonal Antibodies, we chose to assign adequate bioavailability of greater than or equal to 70 % to 1, and inadequate bioavailability of less than 70% to 0. Using SVM, we could find support vectors that could define the hyperplane and margins that would help predict whether there is adequate bioavailability. We decided to make a predictive model using the three different kernels: linear, RBF, and polynomial, and use K-folds to cross-validate each of these predictive models to see which would give the highest score and therefore be the best kernel to use.

## 3 Drug Overviews

### 3.1 Adalimumab

Humira, also known as Adailumbab, is a monoclonal antibody developed by AbbVie that is used to treat patients with rheumatoid arthritis. It does so by targeting a cytokine molecule known as the necrosis factor-alpha(TNF-*α*) which plays a major role in the inflammation immune responses. To do this specifically, Adailumbab binds to the antigen-binding fragment on the TNF transmembrane. This leads to the blockages of the TNFR1 and TNFR2 signaling pathways. One responsibility of the TNFR 1 pathway is the regulation of cytokines that promote inflammatory response and activation of apoptotic signals resulting in cell death [2]. The TNFR2 signaling pathway is responsible for cell activation, proliferation, and helps activate TNFR1 cell death process. Humira has been shown to reduce serum levels of matrix metalloproteinases, MMP-1 and MMP-3, which play a large role in impaired tissue remodeling and repair. A decrease in these proteins will result in lower inflammation. When administering Humira, for a single 40 mg dose delivered subcutaneously, the maximum serum concentration was 4.7 ± 1.6 *μ*g/mL[3]. The time to reach peak concentration was 131 ± 56 hours [3]. The average bioavailability was found to be 64%[3]. In terms of market share, for 20 years, AbbVie had full control of the whole market, generating over 200 billion dollars in sales[4]. In 2023, Amjevita launched a biosimilar with a 5% discount compared to Humira. Another biosimilar by SmithRx, Yumisry, was being sold for a 90% discount compared to Humira [4]. The market share for Adailumbab is becoming more competitive with the advent of biosimilars and existing patent expirations.

**Figure 3:**
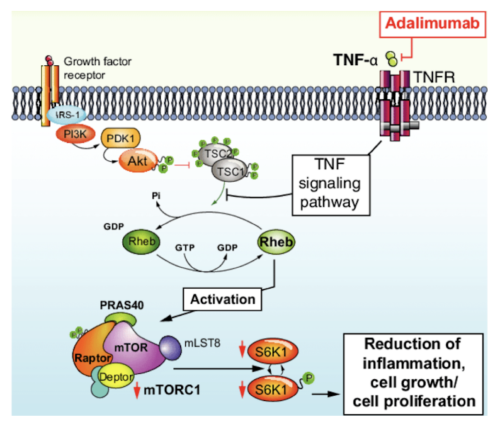
Mechanism of action of Adaluminab binding to the antigen on TNF membrane which ultimately inhibits the TNFR1 and TNFR2 pathways, reducing inflammatory activity characteristic of Rheumatoid Arthritis.[40]

### 3.2 Bevacizumab

Avastin, known as Bevacizumab, is a monoclonal antibody that acts as a vascular endothelial growth factor(VEGF) inhibitor that is used to inhibit angiogenesis. VEGF has been shown to have higher expression levels in solid tumor-type cancers. By binding to the VEGF-A, it inhibits from binding to the endothelial cell surface receptors VEGFR-1 VEGFR-2 which are the signal pathways for new microvascular blood vessels to be formed as well as from tumor blood vessels from growing [6], allowing for a more effective chemotherapeutic delivery. Bevacizumab targets angiogenesis-driven cancers such as non-small cell lung cancer, metastatic breast cancer, renal cell carcinoma, colorectal cancer etc.. When administered subcutaneously, Bevacizumab’s mean concentration-time was 17.38 ug/mL and reached its peak concentration time at about 2.50 hours[7]. Currently, there is no data for the bioavailability of Bevacizmub being delivered subcutaneously which leaves room for further investigations with machine learning predictive models. Since its FDA approval in 2004, Avastin had full control of the market reaching its highest revenue in 2019 at 7.1 billion dollars [8]. As of recently, biosimilars of Bevacizumab released by Amgen called Mvasi in 2017, and Pfizer’s release in 2019 have begun to take a share of Avastin’s market. Combined Am-gen and Pfizer’s Bevacizumab have generated over 2.2 billion dollars in revenue[8]. Because Mvasi is sold at a 23% discount and Pfizer is sold at a 15% discount compared to Avastin, the market share for Avastin is expected to decrease year over year [9].

**Figure 4:**
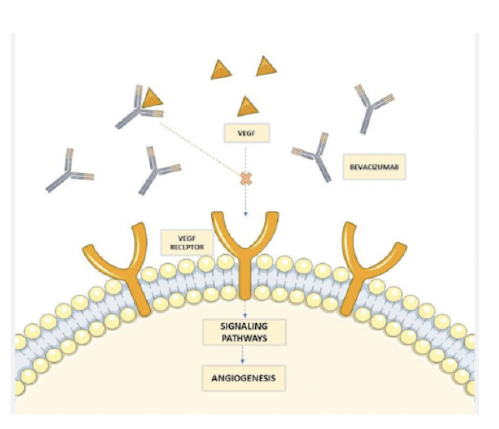
Mechanism of action of Bevacizumab binding to the VEGF-A to prevent angiogenesis from occurring, and preventing the flow of nutrients, oxygen, and growth factors to metastatic cancer growths [41]

### 3.3 Belimumab

Belimumab, known as Benlysta, is a monoclonal antibody that inhibits soluble B lymphocyte stimulator (BLyS) to help treat systemic lupus erythematosus (SLE). BLyS is a cytokine that binds to receptors to help the differentiation of B-cells mature into the antibody-secreting plasma cells [10]. It is also responsible for helping prevent apoptosis. Belimumab can bind to BLyS because Belimumab’s structural properties have a high affinity to the cytokine molecule. By binding to BLyS and blocking the pathway, it reduces the amount of antibody-secreting plasma cells from forming and more autoreactive B cell deaths occur [10]. People with SLE often experience high inflammation and tissue damage. By increasing the amount of autoreactive B cell death, it reduces the onset of inflammatory responses. The pharmacodynamics of Belimumab have shown a decrease in B cells, balanced in IgG levels and anti-dsDNA antibodies [11]. These are all indicators of an appropriate response to SLE patients. The pharmacokinetics in Belimumab show a linear pattern across different doses with maximum concentration for subcutaneous delivery occurring after multiple days [11]. The average bioavailability for Belimumab is 74% [10]. Benlysta was approved by the FDA in 2011 and had exclusivity on the SLE market. Up until 2022, it has generated over $492.2 million in revenue [12]. In 2021, AstraZeneca released Saphnelo as another drug in the SLE market, which is expected to take a part of the market share from Benlysta [12]. It is important to note that Saphnelo uses a different monoclonal antibody, Anifrolumab, which blocks the activity of type 1 interferons to help treat SLE.

**Figure 5:**
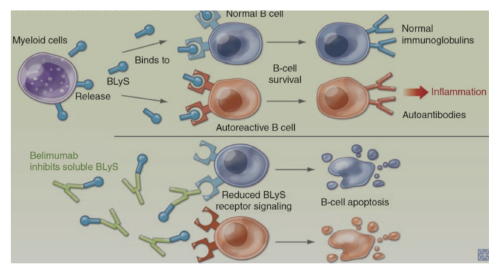
Mechanism of action of Belimumab linking to Blys to inhibit BLyS receptor signaling and therefore promoting cell apoptosis - inducing cancer death [42]

### 3.4 Nivolumab

Opdivo, also known as Nivolumab, is a monoclonal antibody that is used to treat advanced melanoma. By binding to the protein PD-1, it effectively blocks it from binding to the receptor PD-L1 and PD-L2 [13]. The role of PD-1 is that upon binding to PD-L1 it limits the T-cell activation and differentiation to prevent excessive immune responses. In cancers, like advanced melanoma, the PD1-PDL-1 pathway is altered to where there is no immune response. By binding Nivolumab with PD-1, it blocks the pathway for PD-L1 to bind which will create an anti-tumor immune response [13]. The pharmacodynamics of Nivolumab have shown reduced tumor growth in vivo and greater response in T-cells. The inhibited pathway created by Nivolumab also introduced gene expression alterations, osmotic balances, and natural killer cell function[14]. The pharmacokinetics shows a linear pattern across dosages from 0.1 mg/kg to 10 mg/kg that was administered once every two weeks [14]. Although the Nivolumbab is currently administered intravenously (100% bioavailability), subcutaneous delivery is in early-stage trials. The average bioavailability for Nivolumab is still to be determined. Following FDA approval in 2014, Opdivo has shown a steady growth rate with a compound annual growth rate (CAGR) of 6.11 % and reaching over 9.5 billion dollars in sales at the end of 2022 [15]. While there are no biosimilars on the market, Biocad is expected to get FDA approval for its biosimilar to treat metastatic melanoma [15]. For now, Opdivo by Bristol-Myers Squibb Company holds a full share of the market.

### 3.5 Ocrelizumab

Ocrevus, also known as Ocrelizumab, is a humanized monoclonal antibody founded by Genentech that is used to treat multiple sclerosis (MS). It does so by reducing the expression of the cell surface glycosylated phosphoprotein CD20 [16]. CD20 is involved in the development, signaling, and activation of B-cells. By targeting Ocrelizumab to target CD20, a depletion in B-cells occurs. A reduction in B-cells helps reduce the attack on the myelin sheath of the neurons for patients with MS [16]. The pharmacodynamics of Ocrelizumab show a reduction of B-cell count after two weeks of administration into the body. Between each administration, the B-cell count was found to have risen between 0.3% to 4.1%. After about 2.5 years, 90% of patients reached the baseline count for B-cells [17]. The pharmacokinetics for Ocrelizumab show a linear pattern for dosages ranging between 400 mg to 2000 mg [18]. For a 600 mg dose administered every 6 months, the peak concentration is about 212 mcg/mL [18]. In patients with two 300 mg doses administered every two weeks, the maximum concentration reached was 141 mcg/L [18]. Following its FDA approval in 2017, Ocrevus holds one of the highest sales for mAb drugs generating over 5 billion dollars in annual revenue [19]. There are, however, current drugs on the market for MS that are also holding a share of the market. Norvatis released its drug in 2020, Kesimpta, using the monoclonal antibody Ofatumumab which also targets the CD20 protein. It has already generated an annual revenue rate of over 1 billion dollars [20]. Although both are MS drugs, Kesimpta is administered subcutaneously while Ocrevus is administered intravenously. In addition, the price for Ocrevus is about $65,000 a year compared to Kesimpta costing about $83,000 per year [21].

### 3.6 Pertuzumab

PERJETA, also known as Pertuzumab, is a humanized monoclonal antibody that targets the dimerization domain of the HER2 receptor to inhibit the dimerization of HER2 with the other receptors [22]. In normal function, the binding of a ligand to the extracellular domain of the HER receptors triggers a homodimerization or heterodimerization of the HER receptors which triggers a downstream of signaling pathways that are responsible for cell differentiation and proliferation. In breast cancer, unwanted activation of the HER2 receptor caused by tumor cells binding to the receptor can lead to unwanted cell growth and tumor formation. In addition, the different combinations of tumor cells and receptors can lead to a variety of unwanted cellular responses because of the different signaling pathways they can trigger. By Pertuzumab binding to the receptor region of the HER2, it can help prevent tumor growth [23]. It has also been shown that Pertuzumab in combination with Herceptin (trastuzumab) shows an effective combination therapeutic with the death of breast cancer cells [23]. The pharmacodynamics of the HER II receptor have shown in clinical studies of an effective inhibition of the ligand - HER 2 receptor dimerization signaling independent of the HER-2 expression levels [23]. Animal models have also shown that Pertuzumab can be effective for both overexpression and underexpression of HER2 tumors. The pharmacokinetics of Pertuzumab have shown a linear pattern up until a dosage of 25 mg/kg[23]. For studies, using an initial dose of 840 mg followed by a fixed dose of 420 mg every 3 weeks has shown a steady state concentration for tumor growth suppression[23]. Its average bioavailability was 70.1The market share for Perjeta has seen steady growth following its FDA approval in 2012. In 2022, it generated 4.3 billion dollars in sales [25]. While there was a slight decline in Perjeta recently due to the antibody conjugate drug Kadcyla. Both Perjeta and Herceptin, which are both developed by Genentech, continue to dominate the breast cancer drug market [25]. While biosimilars are being developed, Genentech’s drugs remain the leader.

### 3.7 Dupilumab

Dupixent also known as Dupilumab is a monoclonal antibody that is used to treat Atopic Dermatitis (AD). It does this by binding to the IL-4 receptor alpha which blocks the IL-4 and IL-13 pathways because the receptor is not able to dimerize [26]. The IL-4 and IL-13 cytokines are responsible for initiating an immune response such as the development of skin barrier proteins to inflammation and allergic reactions. By binding to the IL-4 receptor it can downregulate the Atopic Dermatitis pathogenesis which would reduce inflammation and improve skin barrier function [26]. Dupilumab’s pharmacodynamic profile normalizes cytochrome P450 levels which may be altered when an IL-4 or IL-13 pathway is activated [27]. Dupilumab is administered subcutaneously with an initial dose of 600 mg. From there, 300 mg doses are administered weekly, and Dupilumab reaches peak concentration one week after the initial dose. The average bioavailability is 64% [27]. Since its FDA approval in 2017, Regeneron Pharmaceuticals and Sanofi Genzyme Dupixent have generated over 10 billion euros in sales [28]. Some competitors are reaching the market such as Leo Pharma’s Adbry recently granted FDA approval and Eli Lilly’s antibody lebrikizumab which was recently approved in the EU but rejected by the FDA in October [29].

**Figure 6:**
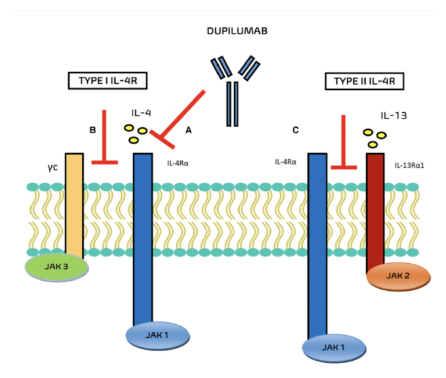
Mechanism of action of Dupilumab binding to the IL-4 alpha receptor to prevent the IL-4 and IL-13 cytokines from binding [46]

### 3.8 Vedolizumab

Entivyo also known as Vedolizumab is a monoclonal antibody that is used to treat ulcerative colitis and Crohn’s disease by binding to the *α*4*β*7 integrin. When the MAdCAM-1 molecule binds to the *α*4*β*7 integrin, it controls the amount of lymphocytes that flow into the gastrointestinal tract and mucosal tissue [30]. When Vedolizumanb binds to the *α*4*β*7 integrin, it stops the migration of the lymphocytes into the gastrointestinal tracts which ultimately reduces the inflammation response[30]. The pharmacodynamics of Entivyo have shown an association between drug concentration and saturation in lymphocytes. This is important for ensuring a proper dosage to achieve maximum benefit. Following an initial subcutaneous injection, dosages of 300 mg are administered every 8 weeks [31]. It has an elimination half-life of about 25 days [30]. The average bioavailability is 80%, a higher average than previously mentioned drugs[31]. In terms of market share, Entivyo made by Takeda has full control of the market as. There are no other biosimilars expected to reach the market by 2032 [32]. Takeda expects to generate over 7.5 to 9 billion dollars annually in the upcoming years [32].

**Figure 7:**
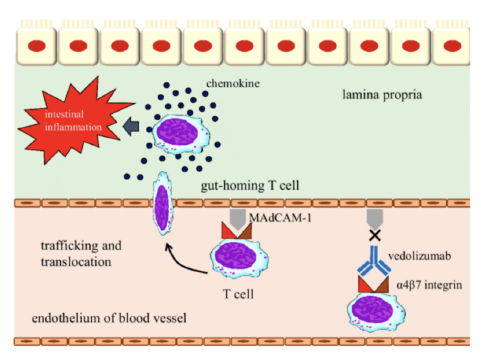
Mechanism of action of Vedolizumab binding to the *α*4*β*7 integrin which prevents the lymphocytes from entering the gastrointestinal tract [47]

### 3.9 Daratumumab

Darzalex, also known as Daratumumab, is a human monoclonal antibody that is used to treat myeloma. It does so by binding to the CD38 glycoprotein which is found on many myeloma cells. CD38 on myeloma cells is responsible for a part of the calcium signaling pathway and metabolism which allows for the proliferation of myeloma cells. Daratumumbab’s ability to link to CD38 effectively induces apoptosis of the myeloma cells and initiates greater T-cell and antitumor activity [33]. It has also been shown that Daratumumab in combination with immunomodulatory agents enhances the activity of T-cells to fight myeloma [33]. Daratumumab in combination with protease inhibitors halt protein degradation and is another effective approach to kill myeloma cells. The pharmacodynamics of Daratumumab have shown a correlation between myeloma protein with a decrease in protein levels with intravenous administration [34]. Upon weekly doses of 16 mg/kg, the maximum drug concentration before the next administration will be about 80% [34]. This shows that drug levels can remain relatively constant. The bioavailability of Daratumumab for subcutaneous administration is 70In terms of the market share for Daratumumab, it is operated by Johnson and Johnson and Genab. Johnson and Johnson originally developed Darzalex. Genab in collaboration with Johnson and Johnson developed a subcutaneous formulation of Daratumumab known as Darzalex FasPro. Johnson and Johnson reported 9.7 billion dollars in sales [36]. Recently, Genab lost a court case to Johnson and Johnson regarding additional royalty payment for Darazlex FasPro in 2030 [36]. While this does not affect the sales now, Gen-mab will not be able to get additional royalties for the new subcutaneous formulation. Despite this, Darazlex FasPro produced about 9.7 million dollars in sales in 2023 [37].

**Figure 8:**
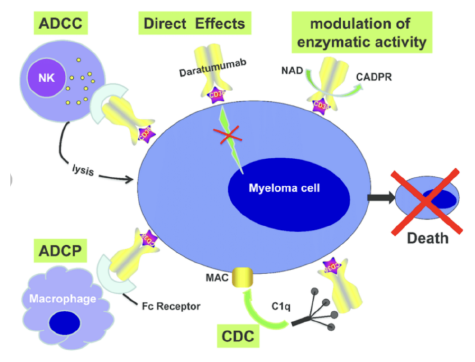
Mechanism of action of Daratumumab with different immunomodulatory agents that ultimately result in the cell death of the Myeloma cell [48]

## 4 Results

As an extension to the model from Han Lou and Michael Hageman, Machine Learning Attempts for Predicting Human Subcutaneous Bioavailability of Monoclonal Antibodies, additional mAb drugs as well as dosage concentration for each mAbs were applied. Below the first two PC components from the PCA analysis are plotted, a heat map that describes how each feature is weighted for each PC component, and lastly the cross-validations scores for when a support vector machine model with a linear kernel, radial basis function kernel, and polynomial kernel were used to predict the bioavailability for monoclonal antibodies.

Based on Figure 9, the monoclonal antibodies are weighted along the PC1 and PC2 axis. There are several of the monoclonal antibodies weighted along the PC2 axis and negatively along the PC1 axis. There is also an outlier, Belimumab along the positive PC1 axis. Because PCA is affected by outliers, this monoclonal antibody could potentially affect which features contribute to the variance of the model - future investigations will mitigate this drug, optimizing the PC analysis resolution.

**Figure 9:**
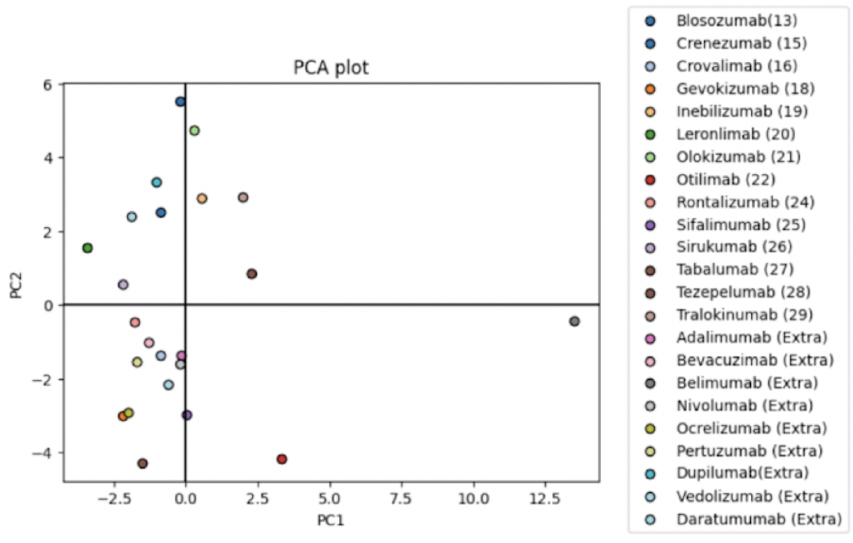
PCA plot of first two principal components showing how each mAb is weighted along the two components that generate the most amount of variance in the model.

Figures 10 and 11 depict the amount of variance explained by the data set. We reduced our 45 features to 10 Principal Components, and Figure 10 shows that these PCs explain about 90 % of the variance in the dataset. In Figure 11, as we increase the principal component the amount of variance decreases. PC1 and PC2 contribute to the majority of the variance as they make up 50% of the variance in the dataset.

**Figure 10:**
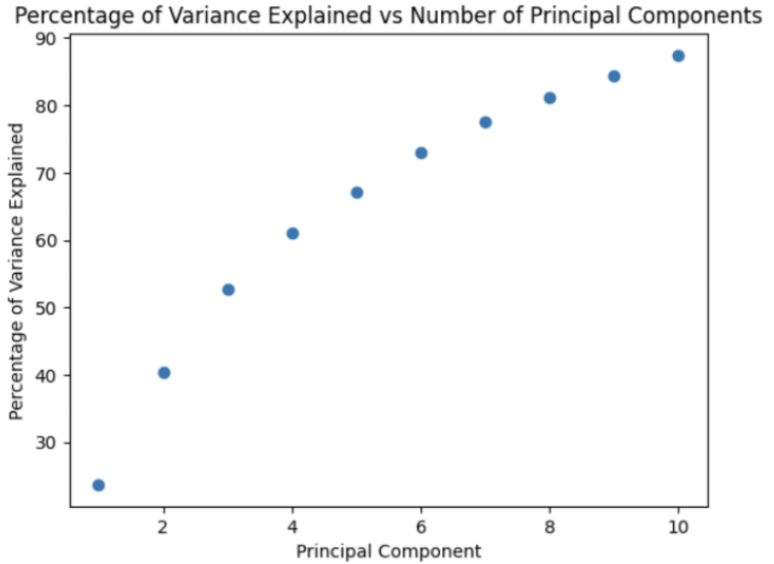
Total amount of variance explained as we increase each component. By the 10th principal component, 90% of variance in the model is explained

**Figure 11:**
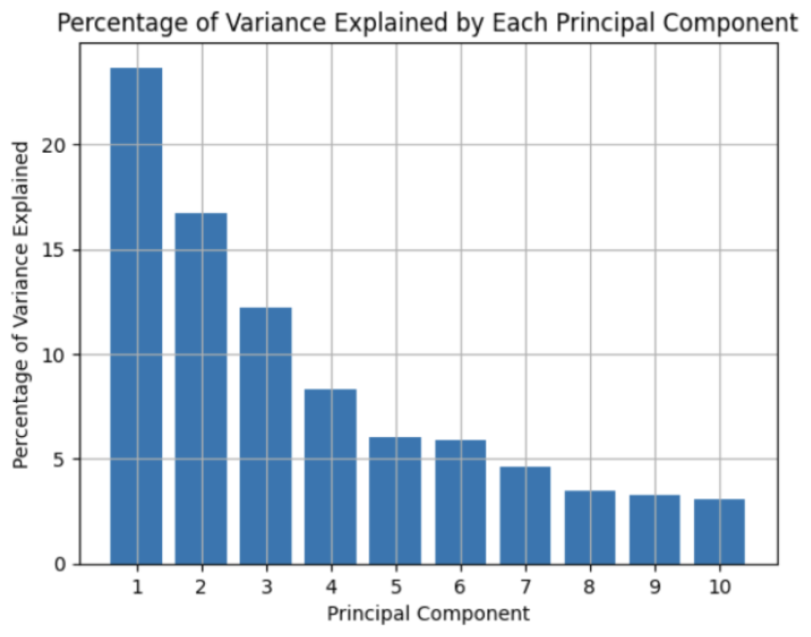
Variance explained by each principal component. As the number of components increases, the amount of variance explained by each component is decreasing.

In Figure 12, each of the features in our dataset contributes to each of the principal components. In PC1, the features of number of amino acids for the heavy chain (−0.27) and total molecular weight (−0.26) make a strong contribution to PC1. In PC2, we see that pI (−0.32) and the charge of the monoclonal body at a pH of 7.4 (0.27) have a strong influence on PC2.

**Figure 12:**
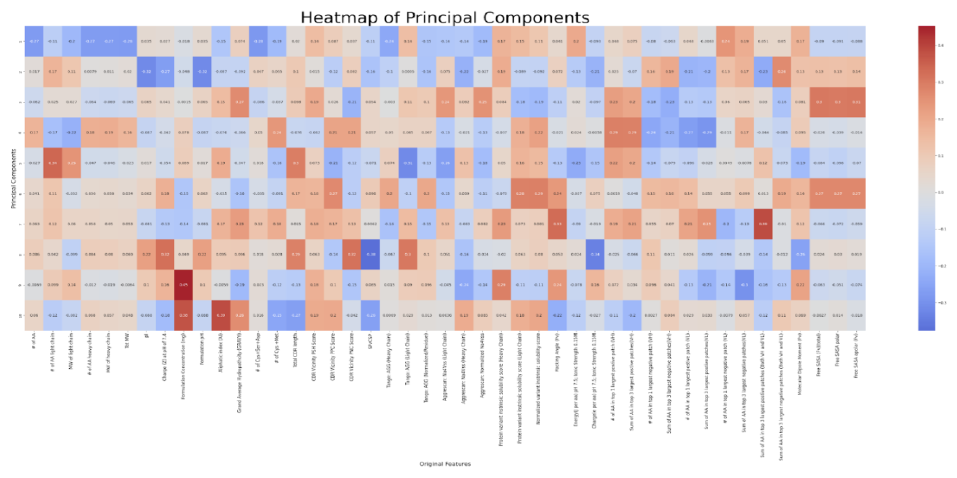
Heat map between the principal components (y-axis) and features (x-axis). Darker and warmer colors represent a strong positive correlation between PC component and feature. Darker and cooler colors represent a strong negative correlation between PC component and feature.

**Figure 13:**
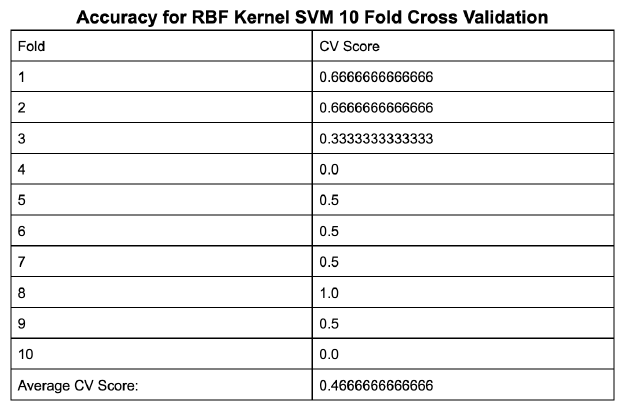
Displays the cross-validation scores for each of the 10 fold for the RBF kernel model as well as the average CV score

**Figure 14:**
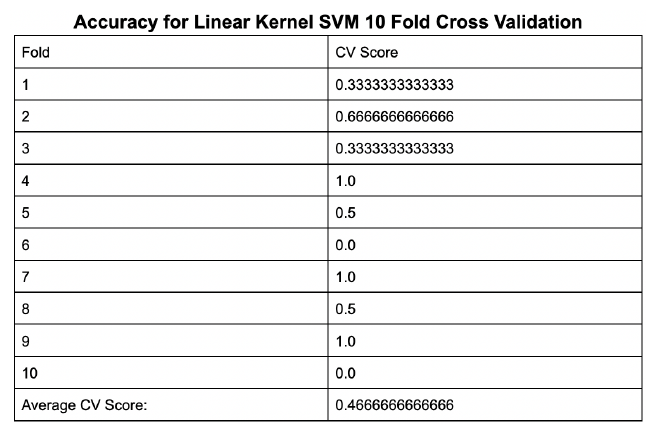
Displays the cross-validation scores for each of the 10 fold for the linear kernel model as well as the average CV score

**Figure 15:**
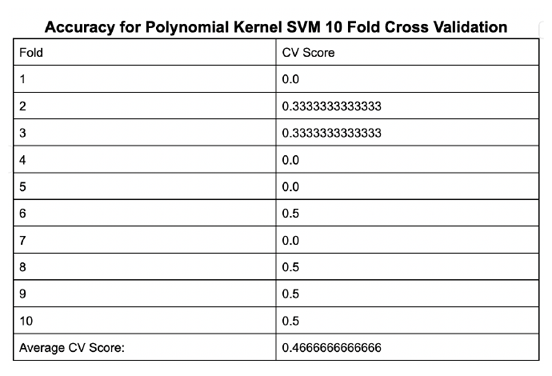
Displays the cross-validation scores for each of the 10 fold for the polynomial kernel model as well as the average CV score

For the first model, an SVM was used with an RBF kernel. Using 10 fold cross validation, for each fold, the dataset was randomly split into 21 samples for the training dataset and 2 samples for the testing dataset. The score helps us determine how well the model predicts whether a mAb has adequate bioavailability. FigureX how it varies for each fold, where sometimes the model predicts the data mostly correctly (score = 0.666), whereas sometimes the model does predict the data right for any instance (score = 0.0). Overall the average score after 10 folds was 0.4666.

For the second model version, an SVM with a linear kernel was used. Similar to the RBF kernel model, a 10 fold CV was implemented, with each fold randomly splitting the training dataset with 21 samples and test dataset with 2 samples. In FigureX, we can see how the scores vary for each fold. Some folds predict the data 100% correctly (score = 1.0) whereas other fold predict it sometimes correctly(score = 0.333). Similar to the RBF model the average score was also 0.4666

For the third model version, a polynomial kernel SVM was used. Using 10 fold cross-validation, for each fold 21 samples were randomly put into the training dataset and 2 samples were split into the testing set. The polynomial SVM has lower accuracy of 0.26666 when predicting if the drug has adequate bioavailability. Currently, the degree of the polynomial is set to the default value of 3, but it will be interesting to see how the model score changes if we change the degree of the polynomial for our model in the future.

Overall, each of the three predictive model methods showed less than 50% accuracy. Several limitations of our dataset could have contributed. Although we collected lots of feature data for this model, ultimately many more samples of monoclonal antibodies are needed for accuracy of prediction. With more data, the outlier that we determined in the PCA plot would have less of an effect on the model. In addition, for the SVM model, we set the gamma and regularization parameter C to their default values. If we manipulate gamma, which controls the significance of a support vector, and the C parameter which is the bias-variance tradeoff, we may obtain a better predictive model.

## 5 Discussion

Based on the drug overviews, mAbs have been able to be used to treat a wide variety of diseases. Whether it’s stopping signaling pathways by binding to cell surface receptors like Humira and Avastin or binding to protein molecules to alter function like Benlysta, Opdivo, and Darzalex, the engineering of mAbs is designed to have a high affinity to these molecules to help treat diseases. With proven effectiveness, the next steps are to continue to find ways to implement subcutaneous formulations. Subcutaneous delivery has been shown to have multiple advantages over intravenous administration. Subcutaneous administration has been shown to reduce the drug burden and patients have been more content with this delivery because of the convenience of administering this at home which would also reduce cost and resources spent [38]. The market analysis for the mAbs shows very little competition among similar formulations.

For example, drugs like Benylysta and Spahnelo are the only two mAbs treating multiple sclerosis. Combined they are holding the majority of the share of the market. Another example is for the treatment of ulcerative colitis and Crohn’s disease. Entivyo has full control of the market for mAb treatment. These examples show the potential drawbacks of few companies monopolizing drugs on the market for diseases as they can control the price of the drug which is often quite costly. To help prevent this, more biosimilars could be generated into the market to facilitate a greater competitive landscape for mAb drugs. This has been proven for mAbs used to stop angiogenesis, like Avastin. Initially, Avastin had full control of the market and was able to set an inaccessible price point.But after biosimilars like Amgen’s Mvasi and Pfizer’s Bevascuzimab were introduced, the market share for Humira declined because the Mvasi and Bevascuzimab were sold at a discount - fostering a greater competitive environment. The current market share described in the drug overview further motivates the need to find a way to help biosimilar generation in the market to distribute the market share and prevent one company from monopolizing the market. Bioavailability is a major factor when assessing the pharmacokinetics of a drug. The amount of drug dosage is inversely proportional to the bioavailability [39]. The greater the bioavailability the less formulation concentration is needed for the drug. This is an important factor if we want to reduce the potency and amount of drugs entering the body. From our drug overview, Adalimumab and Dupilumab showed low bioavailability while monoclonal antibodies such as Bevazcuzimab and Nivolumab have limited bioavailability data. Results like these motivate us to continue to find ways to improve bioavailability for mAbs and to provide a precursor for what the bioavailability could be before clinical trials occur for new mAbs going in the market. In the eyes of the FDA, a drug must exhibit proper biocompatibility or high levels of safety and efficacy to be approved for its intended indications. A low bioavailability reduces the efficacy of the drug therefore affecting its FDA profile; something potentially holding back several drugs from market entry. Models that can predict bioavailability and provide possible explanations for ways to help improve their bioavailability can also speed up the drug clearance process. This is especially important for drugs that have been discontinued such as Rontalizumab (44% bioavailability), Otilimab (35% bioavailability) as well as drugs that are wanting to enter the market.

## 6 Conclusion/Future Perspec tives

This review article aimed to delve into ten monoclonal antibodies (mAbs) and their mechanisms of action, specifically focusing on their pharmacodynamic (PD) and pharmacokinetic (PK) properties. Employing a machine learning model, we endeavor to assess whether these mAbs exhibit adequate bioavailability when administered subcutaneously. Our approach builds off of previous works and by extending this framework, we incorporate an additional set of 10 subcutaneously delivered mAbs, along with considerations of their dosage concentrations (PK property), developing a more sophisticated model. Monoclonal antibodies find application in treating various diseases such as chronic inflammatory conditions, angiogenesis inhibition in cancers, systemic lupus erythematosus, melanoma, multiple sclerosis, breast cancer, and Crohn’s disease, among others. This broad spectrum of applications has led to rapid growth in the immunotherapy field, with an anticipated market size of $679 billion by 2033. Subcutaneous delivery offers numerous advantages over intravenous administration, including reduced drug burden and increased patient satisfaction due to the convenience of at-home administration, which also reduces costs and resource utilization. Models capable of predicting bioavailability and offering insights into improving it can expedite drug clearance processes, a crucial aspect for discontinued drugs such as Rontalizumab and Otilimab, as well as for emerging market entrants.

## Contributions

A.K. O.F. A.J C.K. conceived the review article, collected data, organized figures, and performed all meta-analyses of the literature provided in the paper. C.C. and C.K. contributed to the oversight of this article as co-principal investigators, provided edits, and helped write portions of this article.

## Acknowledgments

The authors are grateful to the various research studies, clinical trials, and individuals behind the work of tackling metabolic diseases. The authors would like to extend their gratitude to the entire Research and Development team at Nexabio Venture Solutions for their guidance, support, and contributions to the overall team dynamics.

